# Transcranial in-depth ultrafast ultrasound brain imaging in non-human primates

**DOI:** 10.1101/2025.08.05.667956

**Authors:** Thomas Orset, Julie Royo, Mickael Tanter, Thomas Deffieux, Michel Thiebaut de Schotten, Pierre Pouget

## Abstract

Ultrasound Localization Microscopy (ULM) offers sub-diffraction resolution vascular imaging by tracking individual contrast microbubbles. However, transcranial implementation in non-human primates remains challenging due to the skull’s acoustic attenuation and distortion. This study presents a method for non-invasive transcranial ULM imaging of the anesthetized squirrel monkey through intact skull and skin. We systematically evaluated key ultrasound acquisition parameters and identified an optimal configuration balancing resolution, depth, and vascular representation. Using these refined parameters, we achieved a system resolution of 9.5 µm, measured by Fourier Ring Correlation (FRC), and consistently visualized vessels as small as 23 µm. Full-brain coronal and sagittal acquisitions revealed major vascular landmarks that were reproducible across animals and slices. Importantly, this resolution enables reliable detection of penetrating arterioles and venules, key elements in neurovascular coupling and implicated in various cerebrovascular pathologies. Finally, by tracking microbubbles, we quantified flow velocities from 8.8 to 55 mm/s, corresponding to flow rates ranging from 0.1 to 279 µL/min, demonstrating the technique’s ability to access micro- and macrovascular dynamics. This work establishes a framework for high-resolution, non-invasive brain imaging in non-human primates, and opens promising perspectives for studying cerebral structure, function, and dysfunction in translational neuroscience.

## INTRODUCTION

Multiple preclinical and clinical imaging modalities have been developed to explore brain vasculature and hemodynamics, each offering a tradeoff between imaging depth and spatial resolution. Angiography at the brain scale can be achieved using a contrast agent in X-ray imaging or through Magnetic Resonance Angiography (MRA). Two-photon microscopy enables high-resolution blood flow imaging at the capillary scale. Still, it is limited to depths of approximately 1mm in the pia mater and at the cost of a cranial window for optical access (*1*).

Initially developed for clinical applications, ultrasound imaging has emerged as a powerful neuroimaging modality in research. Functional ultrasound (fUS), in particular, provides access to vascular anatomy and cerebral blood volume (CBV) changes (*2*).

However, in large animals and primates in particular, ultrasound imaging of the brain has long been restrained by the skull’s aberrating effects, which includes absorption, scattering, and refraction of ultrasonic waves (*3, 4*). In research, these limitations have historically been addressed through invasive approaches such as craniotomy or surgical thinning of the skull, rendering brain ultrasound imaging invasive.

A promising strategy to overcome skull-induced aberrations and provide non-invasive access to brain hemodynamics involves using ultrasound contrast agents. Solutions of injectable microbubbles are used to enhance the blood vessels’ signal for clinical imaging (*5*). Therefore, it compensates for skull attenuation of the signal and preserves imaging depth. Additionally, microbubbles from contrast agents can be used for ultrasound localization microscopy (ULM). Similarly to Photo-activated localization microscopy (PALM), ULM allows one to go beyond the diffraction limit and enables localization with precision superior to the wavelength. A single microbubble can be detected (*6*) and localized with a λ/250 resolution in the axial direction (*7, 8*). It is then used as an individual sensor, and the vasculature is reconstructed by accumulation.

Transcranial imaging of brain vasculature using a microbubble contrast agent has been successfully performed in small animals, such as rats (*9*), and through the temporal window in humans (*10*). Despite being a major challenge, high spatial resolution transcranial ultrasound imaging in non-human primates has still not been done. Squirrel monkeys are widely used for in vivo research in neuroscience. It offers a larger and more folded cortex than the marmoset (*11, 12*), while remaining in the penetration range of high-frequency echographic probes.

Here, we present a method for non-invasive, transcranial ULM imaging of the anesthetized squirrel monkey through intact skull and skin, enabling high resolution imaging across the whole brain, including both superficial and deep vascular structures. We detail our acquisition and image processing pipeline, validate the technique by comparison with other ultrasound imaging studies, and discuss its potential and limitations for further exploration of anatomy and function in the non-human primate brain.

## MATERIAL AND METHODS

### Animals

Recordings were conducted on five adult female squirrel monkeys (Saimiri sciureus) aged 9 to 11 years at the time of acquisitions (two aged 9, one aged 10, and two aged 11). The animals were housed together with their social group. They had free access to water and a balanced *ad libitum* diet composed of pellets, fresh fruits, and food enrichment. Room temperature and humidity were controlled, and the day cycle was 12h long (8:00-20:00). All experiments were ethically approved by the regional ethical committee for animal experiments (Charles Darwin CE005) and the French “Ministère de l’Education, de l’Enseignement Supérieur et de la Recherche” under the project reference APAFIS 21086-202206101449770.

### Acquisition procedure

Animal anesthesia was induced in an induction box with a mix of oxygen and isoflurane (4%). It was maintained with 2% isoflurane using a mask. An intramuscular injection of Dexdomitor (0,007mg/Kg) was made, and a buccal spray of Xylocaïn was applied before intubation. The latter was used to prevent airway obstruction and monitor respiratory function. An infusion was inserted in the lateral tail vein to allow contrast agent injection. Lidocaïne was applied to the animals’ ears before they were placed in a stereotactic frame with ear bars. Body temperature was regulated with heating pads and covers throughout the anesthesia.

Images were acquired with a real-time functional ultrasound scanner prototype (Iconeus and Inserm U1273, Paris, France) and an ultrasonic probe (128 elements, central frequency of 6.25MHz, 250×250µm^²^ of spatial resolution). Centrifuged ultrasonic gel was placed between the probe and the skin.

Each acquisition lasted 3 minutes following the injection of the contrast agent. The contrast agent was Bracco microbubbles (Braco, Italy). A 200µl bolus was injected for each acquisition. It was immediately followed by 200µl of saline to push microbubbles out of dead spaces.

### Ultrasound sequences

To optimize image quality for transcranial ULM in the squirrel monkey, we systematically evaluated the influence of multiple ultrasound acquisition parameters (Table 1). The tested parameters included transmitted frequency (4.167, 5.208, and 6.250 MHz), transmitted voltage (5, 10, 15, 20, and 25 V), number of plane waves used for compounding (5, 7, 11, and 15), and tilt angle (6°, 10°, and 20°). We also examined the impact of frame rate (500 and 1000 Hz), number of half-cycles per pulse (2, 4, 6, and 8), and duty cycle (0.1, 0.25, 0.5, 0.75, and 0.9). The optimal parameter combination, selected based on overall image resolution and vascular detail, consisted of a transmitted frequency of 5 MHz, a transmitted voltage of 5 V, 11 compounded plane waves with a 20° tilt angle, 4 half-cycles per pulse, and a 0.75 duty cycle. A frame rate of 500 Hz was used. These settings were applied for the acquisitions shown in Figures 3, 4 and 5.

**Table 1.** Ultrasound probe parameters. Summary of all tested acquisition parameters. The optimal parameters used for final imaging (figure 3 to 5) are highlighted in bold.

| Parameters | Tested values |
| --- | --- |
| Transmitted frequency (MHz) | 4.167, **5.208**, 6.250 |
| Transmitted voltage(V) | **5**, 10, 15, 20, 25 |
| Number of plane waves | 5, 7, **11**, 15 |
| Tilted angle(°) | 6, 10, **20** |
| Frame rate(Hz) | **500**, 1000 |
| Number of half-cycles | 2, **4**, 6, 8 |
| Duty cycle | 0.1, 0.25, 0.5, **0.75**, 0.9 |

### ULM processing

First, to isolate microbubble signals, a bandpass adaptive singular value deconvolution (SVD) filter was applied to the data. effectively suppressing tissue signals and noise. Then, microbubbles were localized and tracked across consecutive images. Finally, images were reconstructed at pixel sizes of 1, 10 and 20 µm. The 1µm resolution was used for unbiased computation of the Fourier Ring Correlation (FRC), the 10µm images were used to ensure appropriate intensity saturation for image processing, and 20µm images were generated solely for optimal visualization in whole-brain figures.

### Fourier Ring Correlation

The limit resolution of imaging was computed with the Fourier Ring Correlation, a standard method to estimate sub-wavelength resolution in optics and ultrasound imaging (*13*). We used a 1×1µm^²^ voxel size to ensure sufficient spatial frequency range. The imaging plane was first divided into two independent datasets. The two sub-images were reconstructed with a 1µm pixel size. The 2D Fourier transforms of these sub-images are computed, and the image is divided into concentric frequency rings. For each ring, the FRC value is calculated, and spatial resolution is estimated by comparing the FRC curve against the 2σ threshold curve. The first intersection between the two curves is interpolated to define the cutoff spatial frequency, and the resolution is the inverse of that frequency.

### Saturation

To assess the saturation of the ULM images by microbubble tracks, density profiles were computed based on 2D density maps. For each map, a depth profile was generated by averaging the intensity of all pixels within each horizontal row, corresponding to a given imaging depth. This provided a one-dimensional representation of microbubble signal distribution across depth. To reduce noise and emphasize underlying trends, the resulting depth profiles were smoothed using a moving average filter with a window size of 500 pixels. These profiles were used to compare the relative completeness of vascular reconstruction across depth for different acquisition or processing parameters. Significantly, for each profile comparison, only one parameter was modulated at a time, while all others were held constant unless otherwise indicated.

### Diameter and blood flow measurement

The vessel diameter was estimated using a density profile extracted along an orthogonal line to the vessel’s axis. The diameter was then defined as the full width at half maximum (FWHM) of this profile.

For velocity and blood flow measurements, a rectangular region of interest was manually selected along the axis of the vessel within the velocity map. A velocity profile was computed by averaging values along the length of the patch. The maximum value of this profile was taken as the peak velocity Vmax, and the mean velocity was approximated as Vmean=Vmax/2, assuming laminar flow under Poiseuille’s law. Blood flow was then calculated using the estimated diameter and the derived mean velocity.

## RESULTS

Figure 2 presents the results of our systematic acquisition parameters optimization for transcranial ultrasound imaging in the squirrel monkey. The first parameter tested was the transmitted frequency (4, 5, and 6 MHz), presented in Figure 2a. The Fourier Ring Correlation (FRC) curves yielded lateral resolution limits of 9.8, 10.98 and 10.86µm, respectively, with no significant differences. However, the density depth profile revealed a sharp signal attenuation at 6 MHz. While 5MHz provides the highest microbubble detection density at superficial and mid depths, 4 MHz offers better penetration and detection of large deep vessels, though with a lower overall detection density and a more scattered representation of small cortical vessels.

**Figure 1.**
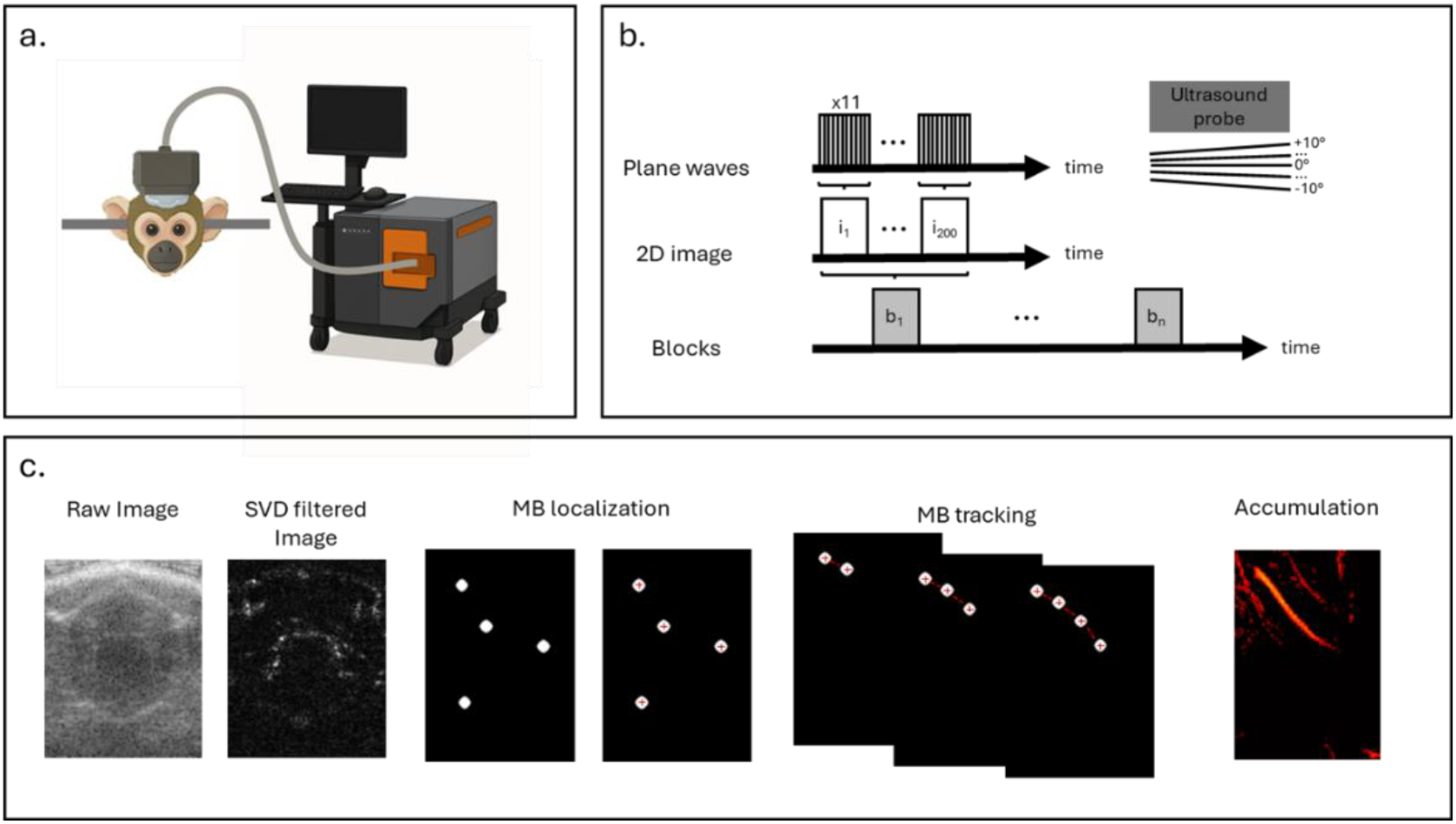
Transcranial Ultrasound Localization Microscopy in the squirrel monkey. a. Acquisition setup: anaesthetized animal is head-fixed in a stereotax. His head is shaved and covered with ultrasonic gel. The IcoDeep ultrasonic probe is placed in the gel and connected to the IconeusOne system. **b**. Imaging sequence: series of plane waves are emitted at different angles and compounded to form a single image. Images are grouped in sets of 200 to form one block. **c**. Image processing pipeline, including SVD filtering, MB localization and tracking (schematic), and accumulation of tracks.

**Figure 2.**
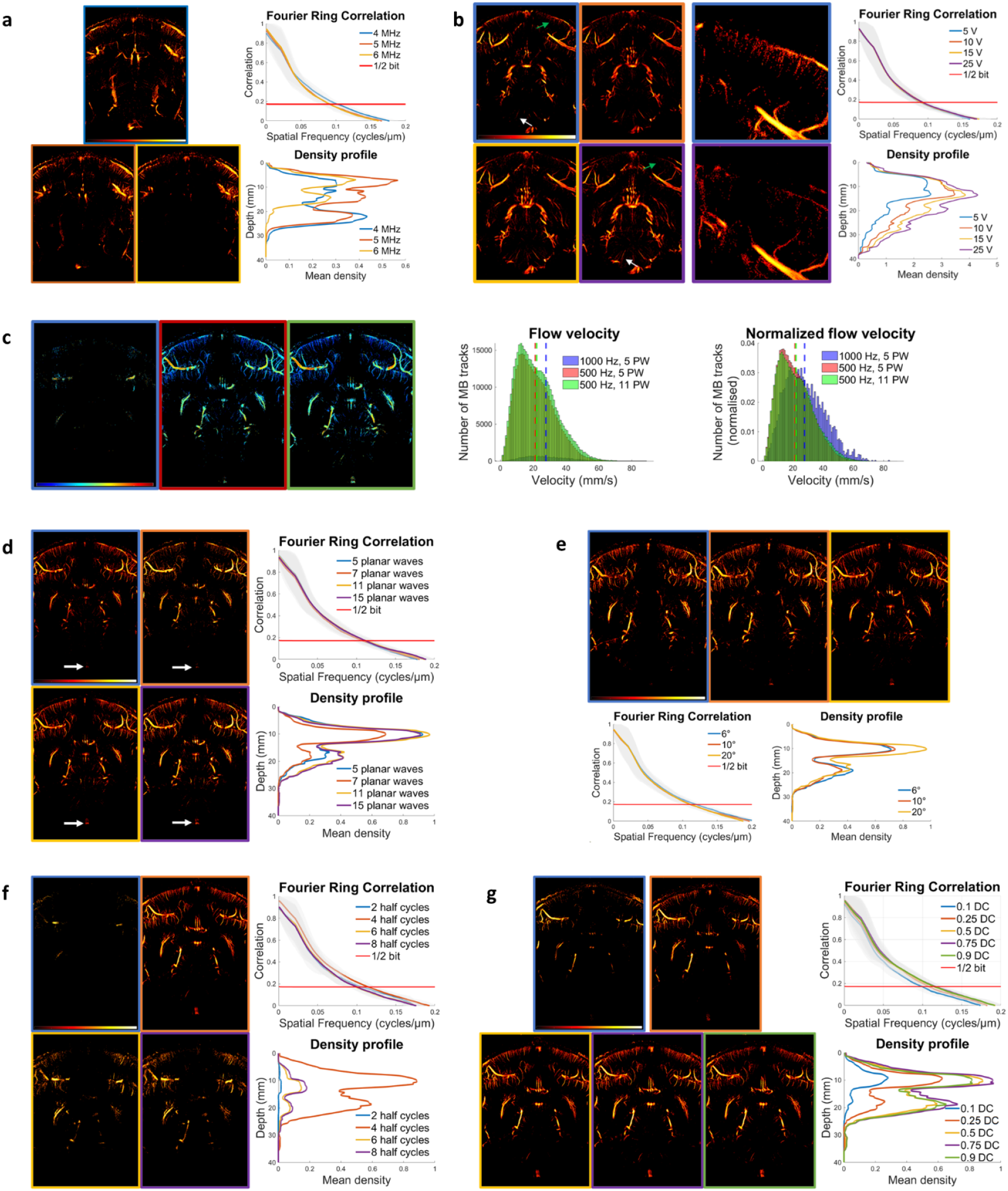
Optimization of ULM acquisition parameters. Images with hot color scale represent microbubble density maps (log scale), ranging from dark red at -30 dB to bright yellow at 0 dB. Images with blue-to-red scale represent velocity maps (0-60 mm/s). Fourier Ring Correlation curves are smoothed and plotted with a shaded envelope indicating fluctuations amplitude. Plots are color-coded to match their corresponding images for each subfigure. **a**. Transmitted frequency: 4, 5, and 6MHz **b**. Transmitted voltage: 5, 10, 15, and 25V. Two enlarged images of the cortical region at 5 and 25 volts are presented in addition to the four complete images. **c**. Frame rate at 1000 and 500(Hz) with 5 or 11PW. In both histograms, dashed lines indicate the mean velocities. **d**. Number of plane waves (PW): 5, 7, 11, and 15. **e**. Angle aperture: 6°, 10°, and 20°. **f**. Number of half-cycles (HC): 2, 4, 6, and 8. **g**. Duty cycle (DC): 0.1, 0.25, 0.5, 0.75, 0.9.

**Figure 3.**
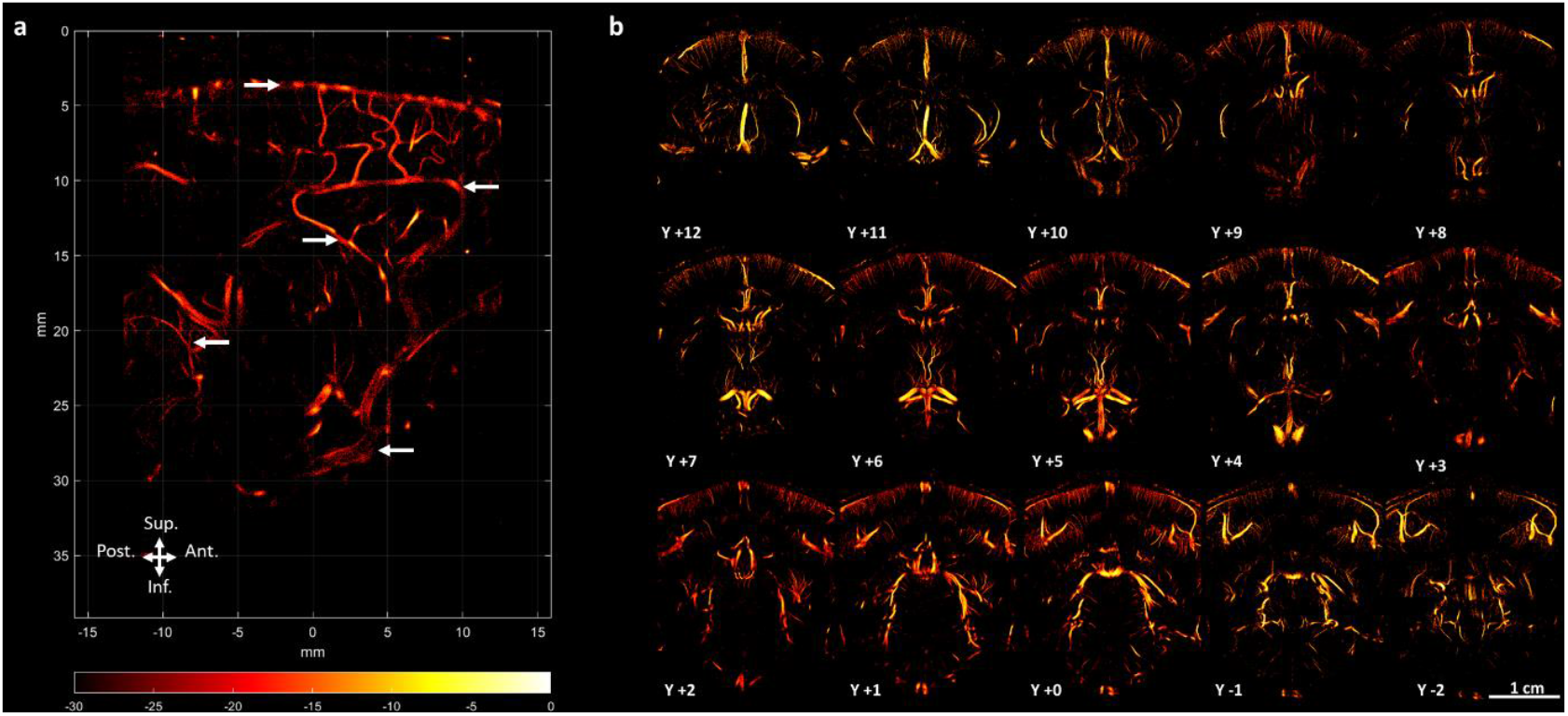
Vascular anatomy of the squirrel monkey. Microbubble density maps (log scale) **a**. Medial sagittal view. Anatomical landmarks (white arrows), from superficial to deep, indicate the Pial Vessels (PV), Pericallosal Artery (PA), Posterior Cerebral Artery (PCA), Superior Cerebellar Artery (SCA), and the Anterior Cerebral Artery (ACA). **b**. Coronal planes acquired every millimeter over 1.4cm. Y0 refers to the ear bar zero reference plane.

**Figure 4.**
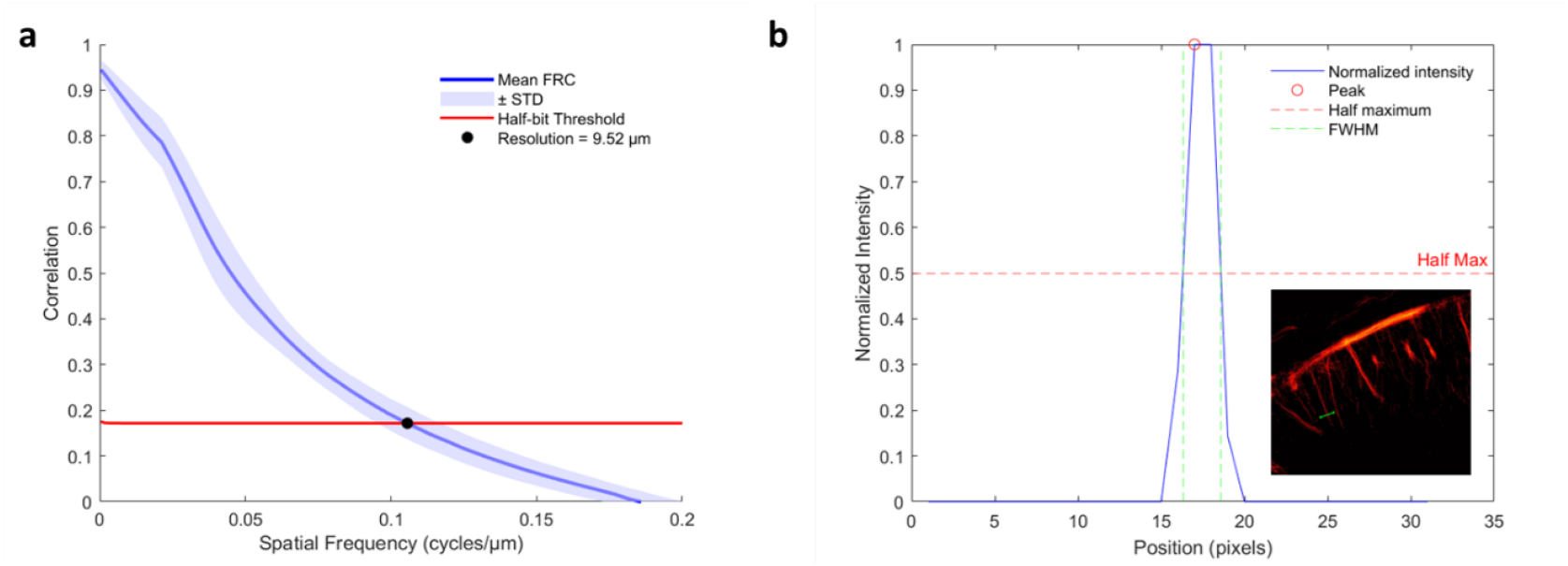
System and image resolution. a. Mean Fourier Ring Correlation (FRC) curve. The solid blue line represents the mean FRC across 24 acquisitions. The shaded envelope indicates the standard deviation. The red curve represents the half-bit threshold. The black dot marks the intersection with the threshold, corresponding to a resolution limit of 9.52µm. **b**. Detectability of subwavelength structure. The solid line corresponds to the normalized intensity profile across a small cortical vessel. Dashed green lines indicate the full width at half maximum (FWHM). The bottom right insert displays the original image with the analyzed vessel section indicated by a green line.

**Figure 5.**
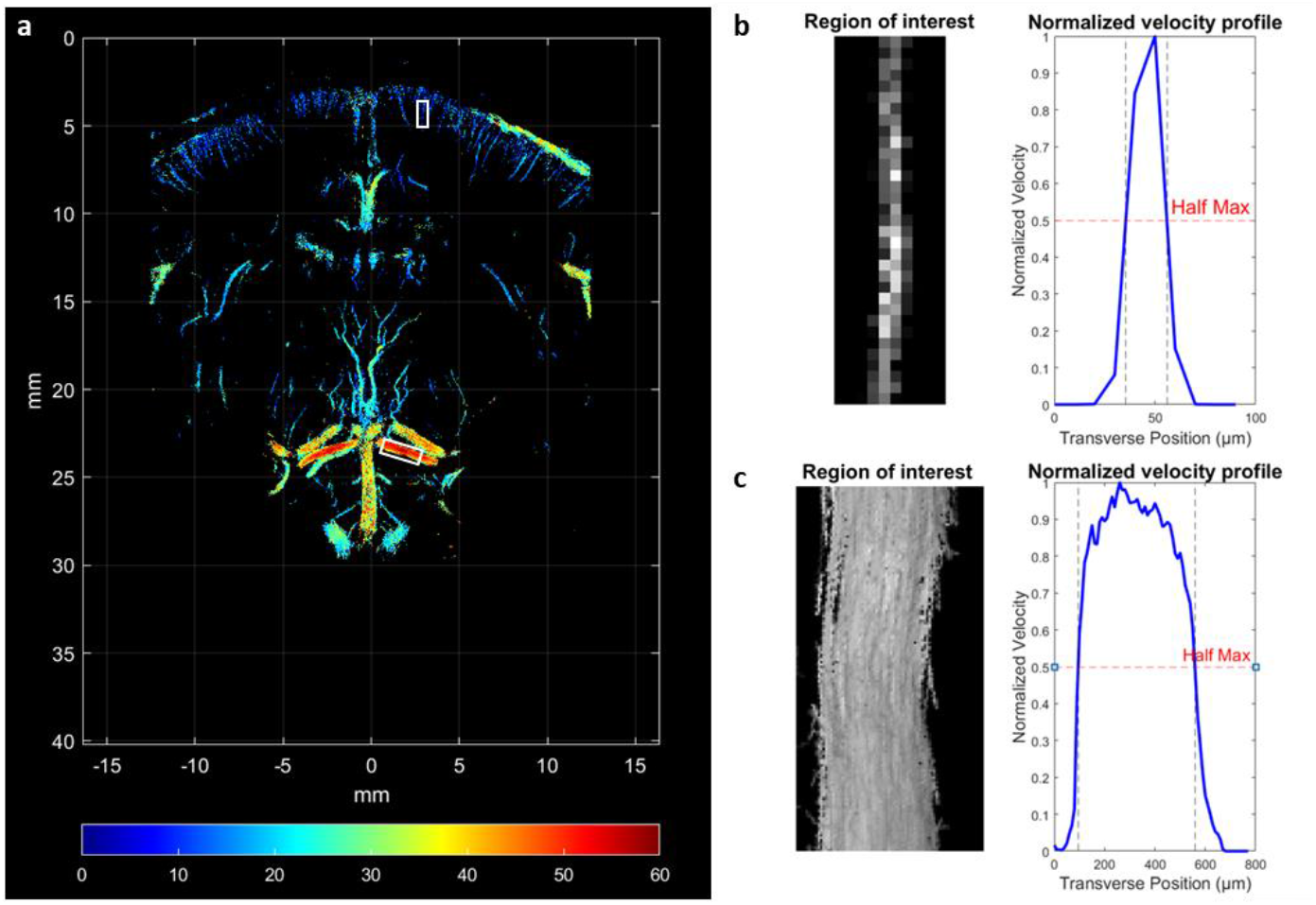
Perfusion properties. a. Velocity map of a coronal plane. The superficial and deep white squares correspond to the regions of interest used in Fig. 2b and Fig. 2c, respectively. **b**. Cortical vessel segment and corresponding velocity profile. Dashed lines indicate the full width at half maximum (FWHM). **c**. Large deep vessel segment and corresponding velocity profile. Dashed lines indicate the FWHM.

The second parameter examined was the transmitted voltage, tested at 5, 10, 15 and 25 volts (Fig. 2b). FRC curves indicate a progressive improvement in resolution with increasing voltage. A similar trend is observed in density profiles, which show higher values and greater depth expansion as the voltage increases. The resulting images provide more details about density repartition. At 5V, density is distributed between large vessels and arterioles, located at superficial and moderate depths. As voltage increases, density becomes increasingly concentrated in large vessels, notably in deeper regions, which are more clearly revealed at higher voltages. On the other hand, the visibility of small cortical vessels gradually decreases with voltage, and they are nearly absent at 25 V (green arrows).

The third parameter was the frame rate, which is constrained by the number of plane waves and is critical for detecting fast flow. We compared acquisitions at 1000Hz and 500Hz frame rates using the same number of plane waves (5), as well as a 500Hz acquisition using 11 plane waves (Fig. 2c). As shown in the normalized histogram, the 1000 Hz condition tended to detect a higher proportion of fast-moving microbubbles, with an average velocity of 24 mm/s compared to 17 mm/s at 500 Hz. On the other hand, the histogram of absolute count shows that the two acquisitions at 500Hz detect significantly more trajectories. This translates as a notably richer vasculature representation in the corresponding images.

We then investigated the effect of angular aperture at +/-6°, 10°, and 20° (Fig. 2d). Images show minimal differences between conditions, and FRC-based resolution estimates are similar. However, the density-depth profiles indicate a higher density of detected microbubbles at ±20°, particularly in deeper regions.

The effect of the number of plane waves (PW) was also assessed for 5, 7, 11, and 15 PW (Fig. 2e). Image analysis reveals improved detection of deep vessels (white arrows) with 11 and 15 plane waves. The density-depth profile is consistent with this observation, showing increased microbubble density at depth. Resolutions measured by FRC are very similar, with 11 planar-waves giving the best resolution (8.94µm) over 15PW (8.97µm).

We next examined the number of half cycles per transmitted pulse, using 2, 4, 6, and 8 half cycles (Fig. 2f). The 2 half-cycle condition results in almost no vascular signal. At the same time, 6 and 8 half-cycles provide a poor signal. The best image quality regarding vasculature completeness is observed at 4 half-cycles. This is confirmed by the density profiles, where 4 half-cycles produce the highest signal across all depths, up to 2.5, while the other conditions remain under 0.5. FRC curves show similar resolution limits across these settings with 4 half cycles giving a slightly better resolution of 8.72µm, compared to 9.46, 10.12, and 9.96µm for 2, 6, and 8 half cycles, respectively.

The last parameter we tested was duty cycles with 0.1, 0.25, 0.5, 0.75 and 0.9 (Fig. 2g). A duty cycle of 0.1 results in an incomplete vasculature representation and 0.25 remains poor. Good vasculature completeness is achieved at 0.5 and above, with the best overall image quality and density at 0.75. Again, FRC curves show comparable resolutions across most conditions, with the best at 0.75 duty cycle (8.56 µm), compared to 10.03, 9.25, 8.72, and 8.67 at 0.1, 0.25, 0.5, and 0.9 duty cycle, respectively.

Figure 3 shows the microbubble density in the medial sagittal plane (Fig. 3a) and in coronal planes (Fig.3b). The medial sagittal view allows for the identification of major vessels, from superficial pial vasculature down to the Circle of Willis. The multiple coronal slices demonstrate the ability to achieve full brain coverage.

Figure 4 provides a detailed assessment of the system and image resolution using the optimized parameters derived from Figure 2: 5 MHz, 5 V, 11 Planar waves, 20°, 4 Half-cycle, and a 0.75 duty cycle. First an average FRC curve computed from 24 images across 2 animals yielded a reliable estimation of the system’s resolution limit, measured at 9.52µm+/-0.93 based on the half-bit threshold (Fig. 4a). Next, an isolated vessel was used to illustrate the method’s ability to resolve small structures in the final reconstructed images (Fig 4b). The full width at half maximum (FWHM) indicated a vessel diameter of 22.8 µm.

Figure 5 presents a velocity map of a coronal plane (Fig. 5a), from which two vessels were selected for case studies. The first selected is a penetrating arteriole located in the cortex (Fig. 5b), with a peak velocity of 8.8mm/s and an estimated diameter of 20.41µm, resulting in a calculated blood flow of 0.1 µL/min. The second vessel is a major deep artery, exhibiting a peak velocity of 55.11mm/s and an estimated diameter of 463.86µm, corresponding to a blood flow of 279µL/min.

## DISCUSSION

Our study presents transcranial ultrasound imaging in the squirrel monkey. We systematically evaluated key ultrasound imaging parameters to optimize image resolution and overall image quality (*14*).

First, consistent with expectation, higher frequencies reduce penetration depth, and we observed a pronounced effect of the transmitted frequency on the imaging depth, especially we observed a sharp signal attenuation at 6MHz (Fig. 2a). We report that 5MHz is the optimal compromise for transcranial imaging in squirrel monkeys as it offers a sufficient imaging depth to observe deep vasculature (3 cm) and sufficiently thin resolution to distinguish small penetrating vessels in the cortex.

Interestingly, we observed that, if higher transmitted voltages (Fig. 2b) provided superior image resolution, greater density, and deeper penetration, it was strongly biased by the large vessels. Indeed, at 25V, main arteries concentrate the signal, and cortical arterioles and venules are nearly absent. This phenomenon may be attributed to the increased susceptibility of microbubbles to disruption under high acoustic pressures, especially in slow-flowing vessels where prolonged ultrasound exposure occurs. At 5 V, although the global density profile is lower, the detection of microbubble tracks is less biased and better distributed across the vasculature, enabling a more accurate representation of both large and small vessels, particularly in the cortex. Therefore, we selected 5 V as the optimal operating voltage, offering the most faithful representation of the fine vasculature.

The number of plane waves and the angle aperture of the compounding sequence only had a limited influence on overall quality and resolution. These parameters have a crucial role in balancing resolution and frame rate. Indeed, the number of emitted plane waves is technically limited by the frame rates. The latter enables the detection of faster movements. However, our results show that a reduction from 1000 to 500 images/second only slightly affects the detected velocities. Still, the increase to 11PW allows a consistent increase in the total number of bubbles that are tracked, and thus the overall quality of the image.

In contrast, parameters such as the number of half-cycles and duty cycle exhibited negligible effect on image resolution and appearance.

Using the optimized set of imaging parameters, we achieved a system resolution of 9.52µm, as estimated by the Fourier Ring Correlation on 24 acquisitions, each from a different slice and across 2 animals. This corresponds to λ/30.

Using the refined imaging parameters, we successfully acquired images across the whole brain of the squirrel monkey, as illustrated in Figure 3. These acquisitions, performed in both coronal and sagittal planes, demonstrate the ability to capture the full depth of the brain, including deep vascular structures such as the Circle of Willis. The images clearly revealed major anatomical landmarks like the anterior cerebral artery, pericallosal artery and superior cerebellar artery, underscoring this method’s structural fidelity and potential.

Multiple acquisitions displayed vasculature with consistent completeness, confirming the robustness and reproducibility of the approach. Furthermore, coronal acquisitions allow for simultaneous visualization of both hemispheres, providing a valuable opportunity to compare homologous regions across sides.

Then, we explored more in detail anatomical images obtained with refined parameters, to assess system and image quality (Fig. 4). We first computed the mean resolution based on 24 FRC curves, each from a different slice and across 2 animals, showing a system resolution of 9.5 µm (Fig.4a). We also demonstrated the possibility of detecting vessels as small as 23µm with a pixel size of 10µm (Fig. 4b). Importantly, this resolution is sufficient to accurately image penetrating arterioles and venules, which are typically over 20µm in the squirrel monkey (*15*). This is particularly relevant for all physiological applications, as parenchymal arterioles represent the primary site of neurovascular regulation through “intrinsic” mechanisms for neurovascular coupling, in contrast with pial arteries on the brain surface, which are regulated exclusively by the sympathetic and parasympathetic inputs (*16*). Additionally, penetrating arterioles and venules dysfunction has been reported to play a key role in diseases like Alzheimer through blood flow reduction (*17*) or laminar stroke when obstructed (*18*). Their structural remodeling after cortical injuries also plays a crucial role in recovering blood flow and behavior (*19*). Therefore, this level of precision is essential for any ULM-based investigation in the squirrel monkey.

To further evaluate the functional possibilities, we explored perfusion properties in the squirrel monkey. Velocity maps (Fig. 5a) generated via microbubbles tracking allowed us to measure blood flow in both a penetrating arteriole (Fig. 5b) and a major deep vessel (Fig. 5c). These two measurements demonstrated flow velocity ranging from 8,8 to 55mm/s, corresponding to a volumetric flow rate of 0.1µL/min and 279 µL/min. This result highlights the technique’s ability to quantify the vasculature dynamics of the squirrel monkey at all scales.

This study opens new avenues for ultrasound imaging in neuroscience by enabling high-definition transcranial imaging, overcoming the long-standing trade-off between invasive techniques and accuracy. By proposing optimized acquisition parameters and leveraging the squirrel monkey model, we identify a promising balance for investigating non-human primates: through the skull and beyond its limitations.

## ACKNOWLEDGMENTS

M.T.d.S is supported by HORIZON-INFRA-2022 SERV (Grant No. 101147319) ‘EBRAINS 2.0: A Research Infrastructure to Advance Neuroscience and Brain Health’, by the European Union’s Horizon 2020 research and innovation program under the European Research Council (ERC) Consolidator grant agreement No. 818521 (DISCONNECTOME), the University of Bordeaux’s IdEx ‘Investments for the Future’ program RRI ‘IMPACT’, and the IHU ‘Precision & Global Vascular Brain Health Institute – VBHI’ funded by the France 2030 initiative (ANR-23-IAHU-0001).

This work is supported by the TOMFU project funded by the Agence Nationale de la Recherche (ANR-22-CE37-0021).

